# Highly accurate estimation of cell type abundance in bulk tissues based on single-cell reference and domain adaptive matching

**DOI:** 10.1101/2023.07.22.550132

**Authors:** Xin-Yang Guo, Zhao-Yang Huang, Fen Ju, Chen-Guang Zhao, Liang Yu

## Abstract

Accurately identifying the cellular composition of complex tissues is critical for understanding disease pathogenesis, early diagnosis, and prevention. However, current methods for deconvoluting bulk RNA sequencing (RNA-seq) typically rely on matched single-cell RNA sequencing (scRNA-seq) as a reference, which can be limiting due to differences in sequencing distribution and the potential for invalid information from single-cell references. To overcome these limitations, we introduced SCROAM, a novel computational method that overcomes these challenges. SCROAM transforms scRNA-seq and bulk RNA-seq into a shared feature space, effectively eliminating distributional differences in the latent space. We then generate cell-type-specific expression matrices from scRNA-seq, enabling accurate identification of cell types in bulk tissues. We evaluated the performance of SCROAM by benchmarking it against simulated datasets and human breast cancer and peripheral blood datasets, demonstrating its accuracy and robustness. To further validate SCROAM’s performance, we conducted single-cell and bulk RNA-seq experiments on mouse spinal cord tissue and applied SCROAM to identify bulk tissue cell types. Our results indicate that SCROAM is a highly effective tool for identifying similar cell types, surpassing the performance of existing methods. We then performed an integrated analysis of liver cancer and primary glioblastoma to investigate the relationship between cell type composition and clinical outcomes in various tumor types, highlighting the significance of SCROAM for understanding cellular heterogeneity in complex diseases. Overall, our work presents a novel perspective to accurately infer cellular composition and expression in bulk RNA-seq, offering valuable insights into disease pathogenesis and potential therapeutic strategies.

## Introduction

Gene expression profiling provides a powerful approach for generating hypotheses about the underlying biological mechanisms that give rise to changes in gene expression under different conditions. This is a critical step in advancing our understanding of complex biological systems and diseases. With over a decade of development, bulk RNA sequencing (bulk RNA-seq) has facilitated the creation of extensive databases, such as the Genotype Tissue Expression Project (GTEx) (Carithers and Moore 2015; Tomczak et al. 2015) and the Cancer Gene Expression Atlas (TCGA) (Tomczak et al. 2015), offering a valuable resource for investigating gene expression in diverse tissues and diseases. These large-scale datasets enable researchers to explore the complex regulatory networks and molecular pathways underlying various biological processes, providing a foundation for the development of novel therapeutic interventions.

However, bulk RNA-seq is commonly limited by cellular heterogeneity, which is a significant challenge across samples. The mixed cell population within each sample results in an overall signature that represents only the average state of the population, thereby limiting the ability to capture the full spectrum of cellular heterogeneity. Single-cell RNA sequencing (scRNA-seq) provides a means of obtaining transcriptomes that are specific to individual cells (Saliba et al. 2014); However, it is often more expensive and technically challenging than mature bulk RNA-seq (Denisenko et al. 2020; Kuksin et al. 2021), which are still widely used due to their cost-effectiveness and scalability. As an alternative, we propose algorithms for computational deconvolution, which is the process of separating a heterogeneous mixture of signals into its constituent cell type-specific signals derived from the data (Avila Cobos et al. 2018; Vallania et al. 2018; Sturm et al. 2019). This approach provides a cost-effective method to obtain cell composition information and significantly increases the speed and scale of related applications, which can be employed in various biological and clinical research contexts. Its value extends to the analysis of specific drug treatment effects or changes in conditions on cell types, providing a highly cost-effective solution.

Many computational deconvolution methods have been devised to identify the composition of cell types or their specific states within bulk samples (Jin and Liu 2021). These methods can be classified into unsupervised and supervised categories depending on whether reference information is available, such as pure cell type expression profiles or lists of marker genes. Fully unsupervised methods that utilize non-negative matrix factorization (NMF) (Avila Cobos et al. 2018) have low deconvolution accuracy and require meaningful gene signatures or expression profiles of distinct cell types for interpretation. Thus, the most frequently employed methods belong to the supervised category, utilizing marker gene-based approaches to quantify each cell type independently in heterogeneous samples by using expression values of specific marker genes. MCPcounter (Becht et al. 2016) aggregates them into abundance scores, and xCell (Aran et al. 2017) performs statistical tests for marker gene enrichment. Other methods, such as DSA (Zhong et al. 2013), MMAD (Liebner et al. 2014) and CAMmarker (Chen 2019), use marker gene lists to guide deconvolution analysis. UNDO (Wang et al. 2015) is an algorithm that automatically detects cell-specific marker genes within the mixed gene expression scatter radius. It then estimates the proportion of cell types in each sample and deconvolutes the mixed expression by generating a cell-specific expression profile. TIMER (Li et al. 2016) is a web-based tool that utilizes a novel statistical method to estimate the relative abundance of six different immune cell types present in the tumor microenvironment. Alternatively, single-cell reference-based methods express the deconvolution problem as a set of equations that represent the gene expression of a sample as a weighted sum of mixed cell type expression profiles. (Finotello and Trajanoski 2018), this approach enables the inference of bulk cell type proportions within a sample using scRNA-seq. Scaden (Menden et al. 2020) is a deconvolution approach that employs a neural network and it trained on large-scale scRNA-Seq datasets to improve robustness. Bisque (Jew et al. 2020) learns bulk gene-specific transformations from scRNA-seq, this approach eliminates the technical bias that may arise due to differences in sequencing techniques between single-cell and bulk RNA-seq. MuSiC (Wang et al. 2019) proposed a weighted non-negative least squares regression framework. This framework simultaneously weights each gene by cross-sample and cross-cell variation. SCDC (Dong et al. 2021) extends the MuSiC method by proposing an ensemble framework that utilizes multiple scRNA-seq datasets as reference and can implicitly address batching effects between reference datasets in different experiments.

In summary, computational deconvolution provides a valuable method to obtain cell composition information and analyze the specific effects of drug treatments or changes in conditions on cell types. However, existing methods have limitations, including their reliance on specific datasets, the need for a user-supplied matrix of cell-type characteristics, and challenges posed by differences in data distribution between single-cell and bulk RNA-seq data. To overcome these limitations, we propose a novel approach that resolves differences in data distribution between single-cell and bulk RNA-seq data. It constructs cell type-specific expression profiles based on scRNA-seq, and importantly, does not require a user-supplied matrix. Our model accurately identifies bulk tissue cell types, addressing the limitations of existing methods and providing a valuable tool for analyzing the effects of drug treatments or changes in conditions on cell types.

## Results

### Method overview

The SCROAM model was developed to estimate cell type proportion in bulk tissues and apply it to study changes in cell type proportions in diseased tissues, providing valuable information for subsequent treatment of the disease. As depicted in Figure 1, SCROAM begins with a single-cell reference and assumes that each cell has been categorized to a fixed set of cell types. SCROAM can then deconvolve bulk samples from the same tissue and estimate the proportion of different cell type present in the sample. When scRNA-seq data is utilized as a reference for cell type deconvolution, the difference in sequencing distribution from bulk RNA-seq must be taken into account. When constructing a feature matrix from scRNA-seq, SCROAM is not based on the average expression, but instead, each gene is weighted according to its cell-specific score, allowing for larger gene sets to be used in deconvolution. Further details regarding this approach can be found in the Methods section below.

**Figure 1.**
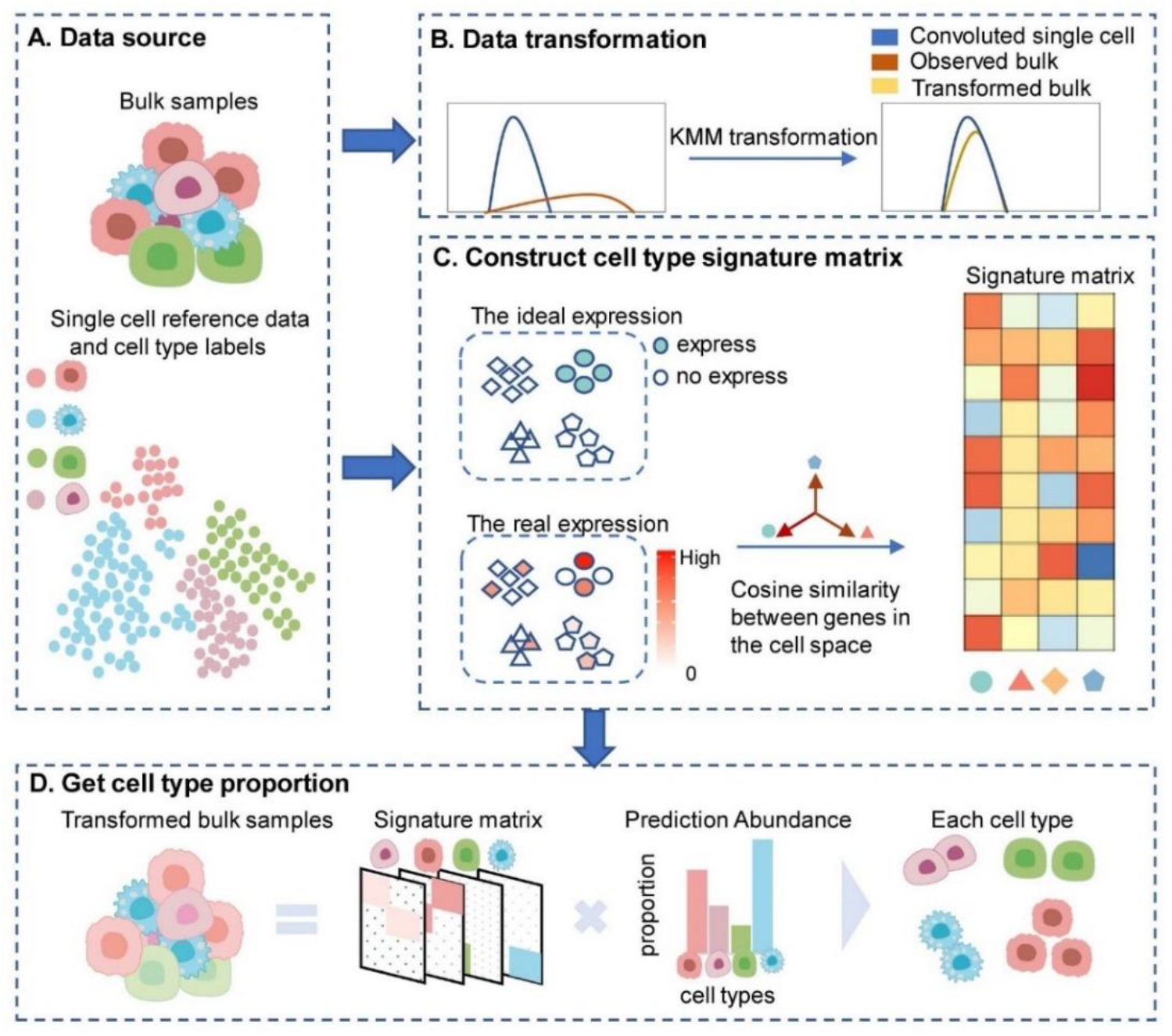
Overview of SCROAM. (a) Our deconvolution model that uses a reference requires two input datasets: bulk RNA-seq count and a reference containing counts of scRNA-seq reads. Additionally, the single-cell transcriptome data must label the cell type to be quantified. (b) SCROAM learns gene-specific transformations of bulk data by utilizing the reference sequences observed in single-cell data. This allows us to account for potential technical bias between sequencing technologies used in single-cell and bulk RNA-seq data. (c) SCROAM begins with scRNA-seq data and classifies the cells into different cell types, which are represented by different colors in the analysis. By calculating gene specificity in a given cell type, an expression matrix reflecting cell type specificity is constructed. (d) SCROAM employs single-cell reference data to estimate the cell type ratio in transformed bulk data.

### Performance on simulated data

To establish the validity of SCROAM, we performed in silico experiments using scRNA-seq data obtained from the Tabula Muris Senis Consortium (2020). We analyzed eight organs, each containing three to nine cell types, and generated “pseudobulks” by pooling reads from known proportions of labeled cells. This allowed us to evaluate the performance of SCROAM in comparison to recently published methods and Non-Negative Least Squares (NNLS). To evaluate the performance of the different methods, we calculated the L1 distance (absolute difference) between the estimated cell type proportions and the true proportions, and then divided the result by the number of cell types present in the sample. We observed that SCROAM had the lowest mean error among the evaluated methods in the reference/bulk configuration. (Fig. 2a, b). Additionally, we visualized the aggregate error by organ (Fig. 2c, d) and found that SCROAM was robust across different contexts. We directly compared the error of each method with that of SCROAM on the same deconvolution task, and found that SCROAM produced less error than the other methods in most cases. Although the nominal improvement of SCROAM in averaging L1 metrics per cell type initially appears small, it is important to note that most tissues are comprised of multiple cell types. Therefore, the overall error can accumulate quickly. Additionally, since the counts for each organ were obtained using both Smart-seq2 and 10x Chromium protocols, cross-protocol comparisons could be easily performed. This is especially important given the different techniques required to generate bulk and single-cell RNA-seq samples. Overall, the findings of this study indicate that SCROAM outperforms other existing methods, particularly in the Smart-seq2 reference and 10x Chromium pseudobulk configuration. Furthermore, the robustness of SCROAM across different organs and experimental protocols highlights its potential as a universal method for deconvolving cell type-specific gene expression from bulk RNA-seq data. For a detailed summary of our results, please refer to Supplemental Table S1 and S2.

**Figure 2.**
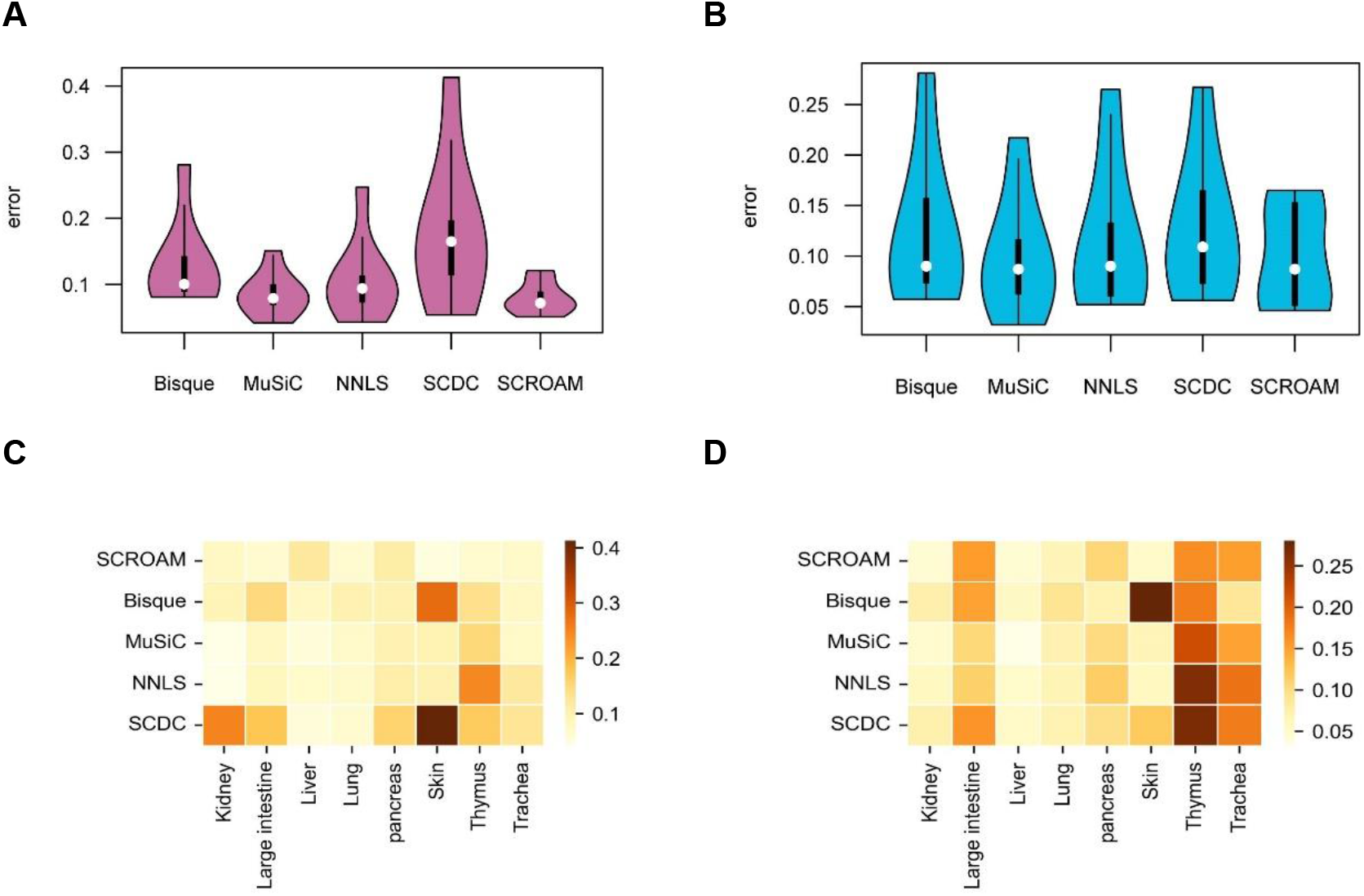
The figure displays the error distribution for each method in the pseudobulk experiment utilizing data from the Tabula Muris Senis dataset. The experiment was conducted on eight distinct organs, and the errors were computed as the mean L1 error across various cell types in each organ. (a, c). show the results for the Smart-seq2 reference and 10x Chromium pseudobulk. (b, d). show the results for the 10x Chromium pseudobulk and Smart-seq2 pseudobulk. In the violin plots, the distribution of errors for each evaluated method is presented, with white dots indicating the mean error. The grid plots use colors to indicate the difference between the mean errors of the different methods in that organ, with darker reds indicating relatively poorer performance. These visualizations allow for easy comparison of the performance of different methods across different organs and experimental conditions.

Given the many factors that need to be considered, including different algorithms, tissues, cell types, and experimental protocols, any benchmark assessment of cell type deconvolution methods must include a large number of combinations. Furthermore, due to the randomness of the data makes it virtually impossible for any one algorithm to outperform the others in all cases, as discussed in Menden (Menden et al. 2020). As a result, we suggest that evaluations should focus on a composite measure of accuracy across multiple cases.

In the data processing step of the SCROAM method, the KMM algorithm is utilized for data transformation. The performance of the deconvolution results can be improved or somewhat reduced depending on the dataset. If the gap between the single-cell reference and bulk data used is significant, the KMM algorithm can greatly enhance the performance. To better understand the effect of the data transformation step, we analyzed and displayed the Large Intestine organ dataset (with Smart-seq2 as the reference). As shown in Fig 3.a, the data transformed by KMM yielded a lower error rate than the data without transformation. Furthermore, as shown in Fig 3.b, we utilized the Jensen-Shannon Divergence (JSD) (Menéndez et al. 1997) indicator to measure the distance between the two distributions. It can be observed that the distance between the bulk data and the cells in the single-cell reference was significantly smaller after data transformation compared to the original bulk data. In Fig 3.c, we analyzed each sample of the Large Intestine organ and assessed the correlation between samples using the Pearson correlation coefficient (PCC)(Cohen et al. 2009). The results showed that applying deconvolution on the Large Intestine data improved the overall evaluation results. These findings suggest that data transformation using the KMM algorithm can significantly improve the performance of SCROAM, particularly when there is a large discrepancy between the single-cell reference and bulk data.

**Figure 3.**
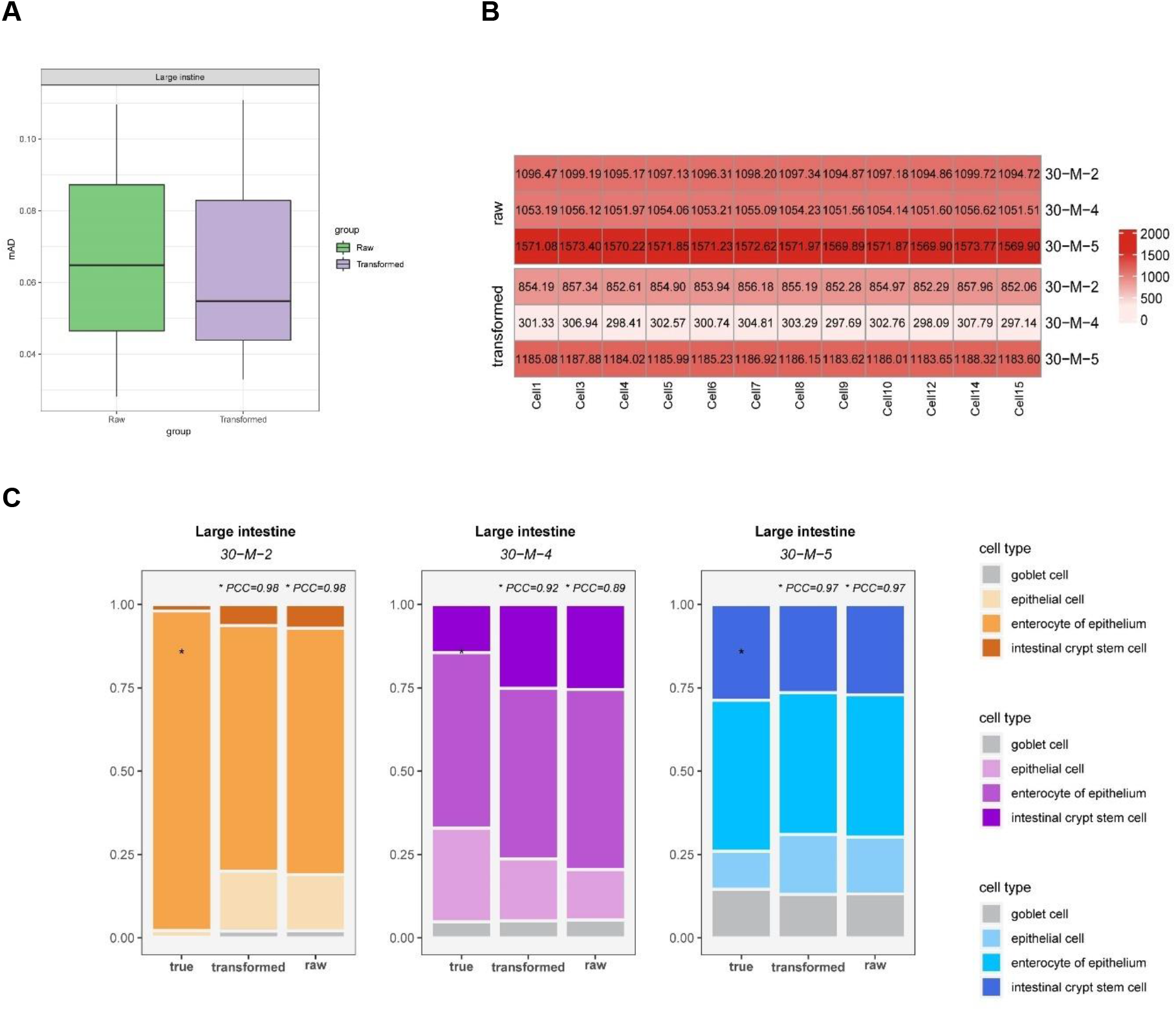
depicts the results of the Large Intestine organ dataset using Smart-seq2 as a reference. (a) The comparison of results before and after data transformation is shown, indicating that the data transformed by KMM resulted in lower error rates. (b) The distance between the raw bulk data and the single-cell reference is compared with the distance between the transformed data and the single-cell reference. The value in each box represents the JSD distance between the sample and the cell. The results show that the distance between the transformed data and single-cell reference is significantly smaller than that of the raw bulk data, highlighting the effectiveness of the KMM data transformation step. (c) The deconvolution analysis results for each sample are presented, demonstrating that the results using transformed data are generally higher than those without transformation.

### Validation with real bulk RNA-seq data

In rare cases, when experimental cell type proportions or bulk RNA-seq samples are available, it becomes feasible to evaluate the real-world performance of cell type deconvolution methods. In this study, we considered three such datasets. The first dataset was derived from the bulk RNA-seq mixture of two human breast cancer cell lines and fibroblasts (60% MDA-MB-468, 30% MCF-7, 10% fibroblasts). To validate our results, we utilized scRNA-seq data published by Dong(Dong et al. 2021) (Fig. 4a). Figure 4b illustrates that SCROAM yielded highly precise results, displaying the lowest mean error out of all evaluated methods. In contrast, the other assessed methods significantly overestimated the bulk fraction of the MDA-MB-468, while underestimating the fraction of the MCF-7 cell line, with the exception of NNLS. These findings suggest that SCROAM is capable of accurately deconvolving cell type-specific gene expression from bulk RNA-seq data, even in challenging real-world scenarios, outperforming other commonly used methods.

**Figure 4.**
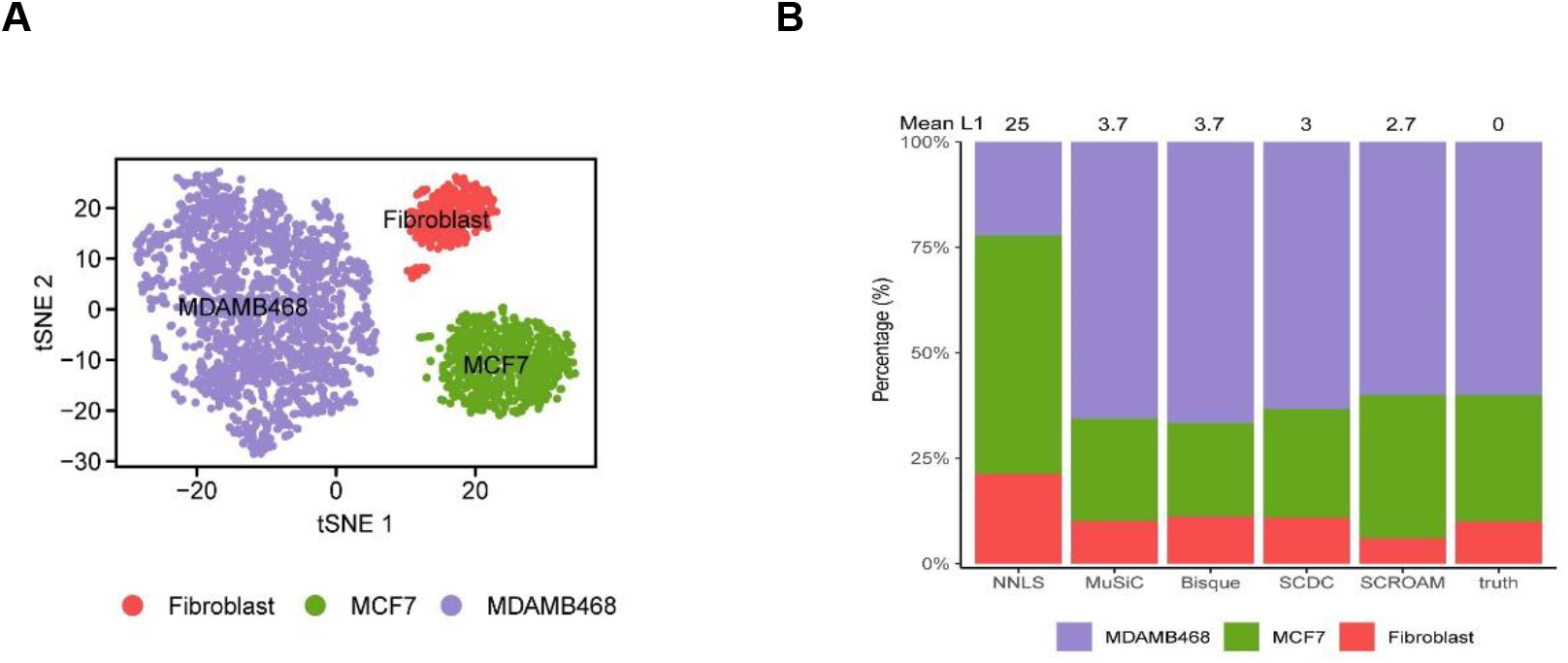
shows the evaluation of each applicable method using data from Dong(Dong et al. 2021), which includes known cell type proportions. (a) shows the single-cell clustering results and t-SNE visualization of the three cell types in the dataset, MDA-MB-468, MCF-7, and normal fibroblasts, with a ratio of approximately 6:3:1. (b) The benchmark of deconvolution results for bulk RNA-seq samples generated by different methods is presented. The proportion estimated by SCROAM has the lowest Mean L1 errors (2.7) to the ground truth, indicating superior accuracy in estimating cell type proportions.

To validate the accuracy of heterogeneous expression measurements in peripheral blood mononuclear cells (PBMCs), we utilized a larger dataset consisting of two sets of 12 bulk whole blood samples from Newman (Newman et al. 2019) and Monaco (Monaco et al. 2019), respectively. The cell types contained in these bulk samples are: B cells, CD4+ T cells, CD8+ T cells, monocytes, natural killer (NK) cells, and neutrophils granulocytes. For deconvolution analysis, we acquired two distinct scRNA-seq PBMC reference datasets. The first dataset was from Newman (Newman et al. 2019) for Newman bulk tissue samples, while the second dataset was a combination of two publicly available reference sets from 10x Genomics for Monaco bulk samples. Notably, since neutrophils are challenging to measure at the single-cell level, resulting in their absence from the majority of the single-cell references. Here we refer to Erdmann-Pham’s (Erdmann-Pham et al. 2021) approach.

To account for their presence in bulk samples, we merged a publicly available scRNA-seq (Xie et al. 2020) dataset containing human neutrophils into the two single-cell references. To account for the limited number of cells in the Newman reference dataset, we undertook subsampling of the neutrophil data to 250 cells. In contrast, we sampled 1250 neutrophils for the reference set in the 10x Genomics PBMC reference dataset.

Our deconvolution analysis showed that SCROAM outperformed all other methods, based on the mean absolute deviation (L1 error) of the two analyses. Table 1 summarizes the results of the analysis. Overall, our findings validate the accuracy of heterogeneous expression measurements in PBMCs and demonstrate the superiority of SCROAM in accurately deconvoluting bulk RNA-seq data into its constituent cell types.

**Table 1.**
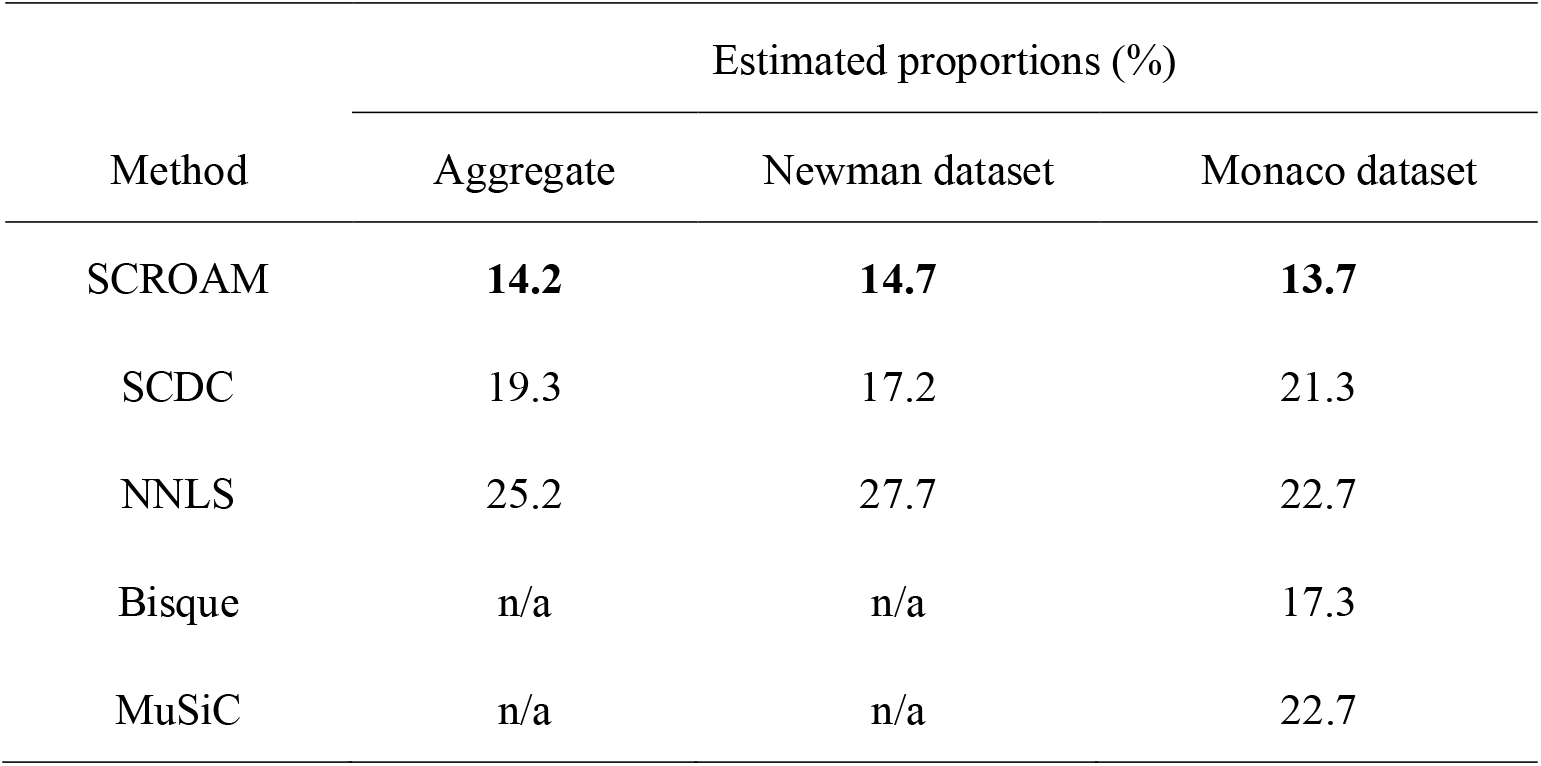
Average L1 errors with PBMC data and ground-truth cell proportions obtained from cytometry.

| Method | Estimated proportions (%) |  |  |
| --- | --- | --- | --- |
|  | Aggregate | Newman dataset | Monaco dataset |
| SCROAM | **14.2** | **14.7** | **13.7** |
| SCDC | 19.3 | 17.2 | 21.3 |
| NNLS | 25.2 | 27.7 | 22.7 |
| Bisque | n/a | n/a | 17.3 |
| MuSiC | n/a | n/a | 22.7 |

The average L1 errors of the two PBMC datasets are presented in the first two columns, while the last column summarizes the L1 errors of the two datasets. The best performing method is indicated in bold. It is worth noting that Bisque and MuSiC were unable to provide scale estimates for the Newman (Newman et al. 2019) data due to the absence of multiple samples of all reference cell types.

### Analysis of mouse Spinal cord injury data

In this study, the actual bulk RNA-seq data acquired from the neural stem region of the mouse spinal cord was analyzed using SCROAM in this study. We first performed scRNA-seq data and bulk RNA-seq data on the identical tissue and used the Seurat (Macosko et al. 2015) pipeline for single-cell clustering, t-SNE visualization of the resulting clusters is presented in Fig. 5a (see the Materials section for details). Through cluster annotation analysis, we identified six distinct cell types: Dormant-Neural Stem Cells (dNSCs), Primed-Neural Stem Cells early (pNSCs_early), Primed-Neural Stem Cells late (pNSCs_late), Active Neural Stem Cells (aNSCs), Transit-Amplifying Progenitors (TAPs), and Neuroblast (NB).

**Figure 5.**
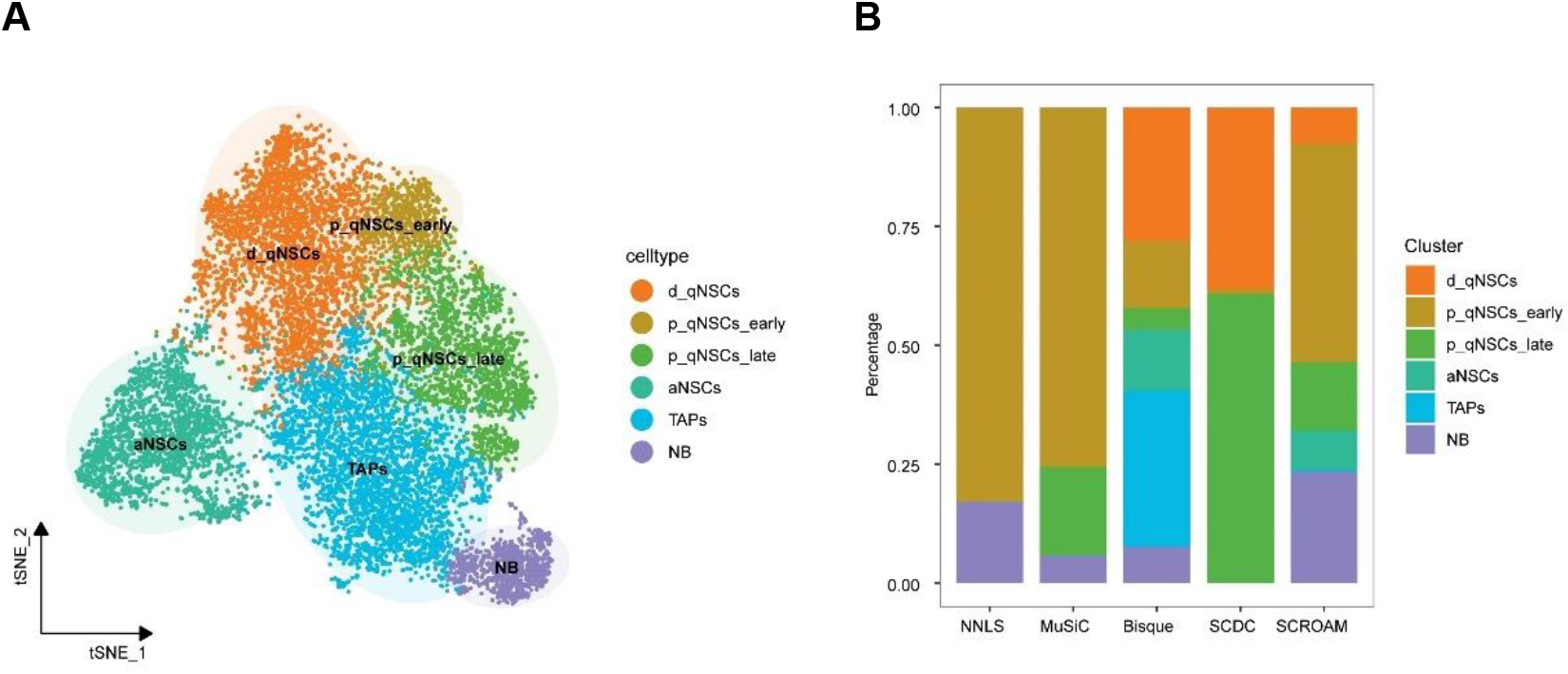
Each applicable method was evaluated using data from the neural stem region of the mouse spinal cord. (a) Following single-cell clustering, t-SNE visualization was generated, revealing six clusters: d_qNSCs, p_qNSCs_early, p_qNSCs_late, aNSCs, TAPs, NB. (b) In the benchmarking of deconvolution results on bulk samples generated by different methods, SCROAM was observed to provide the most accurate estimation of the actual biological proportions among all the benchmarked methods.

Next, we employed annotated single-cell reference data for deconvoluting bulk RNA-seq samples. SCROAM largely follows the w-NNLS framework suggested by MuSiC (Wang et al. 2019), but it varies in several respects. First, SCROAM reduces the difference between single-cell sequencing and bulk sequencing through KMM (Gretton et al. 2009) conversion. Second, when constructing a cell type-specific expression matrix, SCROAM determines specificity by calculating the cosine similarity between genes and calculating gene-specific scores so that genes with smaller scores contribute less to the composition estimate. On this real dataset, we demonstrated that the SCROAM was superior to other methods in accurately determining the proportions of the six cell types present in the bulk sample. (Fig. 5b). The NNLS method only estimated two cell types, while the MuSiC and SCDC methods estimated three cell types. Although the Bisque method was able to deconvolute each cell type, the bulk sample used was from adult normal spinal cord cells, where there are more quiescent cells. The SCROAM approach better reflects this biological reality, demonstrating its utility in accurately deconvoluting similar cell type mixtures from bulk RNA-seq data, emphasizing its utility. These findings highlight the potential of SCROAM as a valuable tool for analyzing bulk RNA-seq data in various biological contexts.

## Applications

### Case study1: Cell Fraction of HCC TME Correlates with Clinical Outcome

The aim of this study was to analyze the tumor microenvironment (TME) in liver cancer by estimating cellular components through transcription using two single-cell RNA-seq reference: (1) GSE115469 (MacParland et al. 2018) (Normal Liver), and (2) GSE146409 (Massalha et al. 2020) (TME-Stroma). GSE115469 represents a liver subgroup in the Human Cell Atlas, while GSE146409 includes the TME of liver cancers, such as Hepatocellular carcinoma (HCC) and cholangiocarcinoma (CCA), and describes over 20 cell types or subtypes. The expression matrices in both datasets are normalized to 10,000 counts/cell and are provided in H5AD file format, along with metadata annotations by the authors for downstream analysis.

Our study aimed to determine if changes in liver cells have an impact on the clinical prognosis of HCC. To achieve this, we conducted survival analysis using the TCGA-LIHC database, the most cited HCC research cohort containing a sizable sample of 370 HCC patients with well-annotated follow-up information. This database is a pooled study of 5 different cohorts with varying risk factors, and it includes survival analyses such as overall survival (OS)(Ally et al. 2017). Among all estimated cell types, Liver sinusoidal endothelial cells (LSECs) and cholangiocytes had a significant impact on patient prognosis. A high fraction of LSEC cells was associated with longer OS (Fig. 6a), while patients with a high proportion of cholangiocytes had a lower OS (Fig. 6b). These findings are consistent with previous reports (Poisson et al. 2017; Chung et al. 2018; Pinto et al. 2018; Gracia-Sancho et al. 2021).

**Figure 6.**
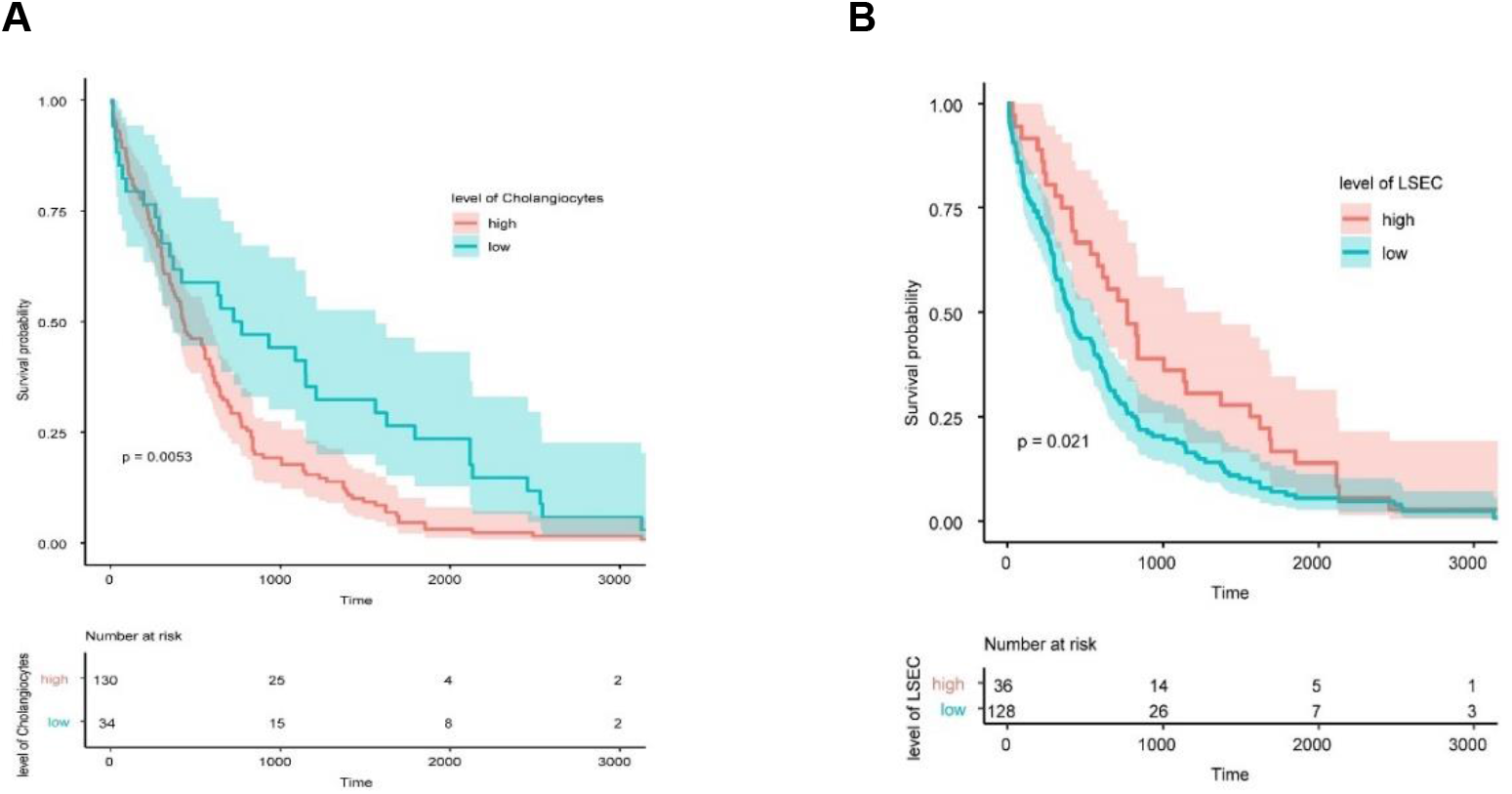
Effect of cell ratio on patient survival. (a) Effect of LSEC cell fraction on overall survival (OS), with patients exhibiting high levels of LSEC cells having longer survival times. (b) The effect of cholangiocyte fraction on OS, with patients having a high proportion of cholangiocytes associated with a lower OS.

In conclusion, our results suggest that both LSECs and cholangiocytes are crucial in determining the clinical prognosis of HCC. The findings of this study have important implications for understanding the biology of liver cancer and developing new therapeutic strategies.

### Case study2: Tumor microenvironment infltration estimation

In this study, we analyzed cell type proportions in 169 glioblastoma (GBM) samples obtained from TCGA(Brennan et al. 2013). To ensure the accuracy of our results, we employed scRNA-seq data obtained from the same tumor type to perform the deconvolution task (Yuan et al. 2018). Using these reference sets, we identified six cell types of GBM (Fig. 7a).

**Figure 7.**
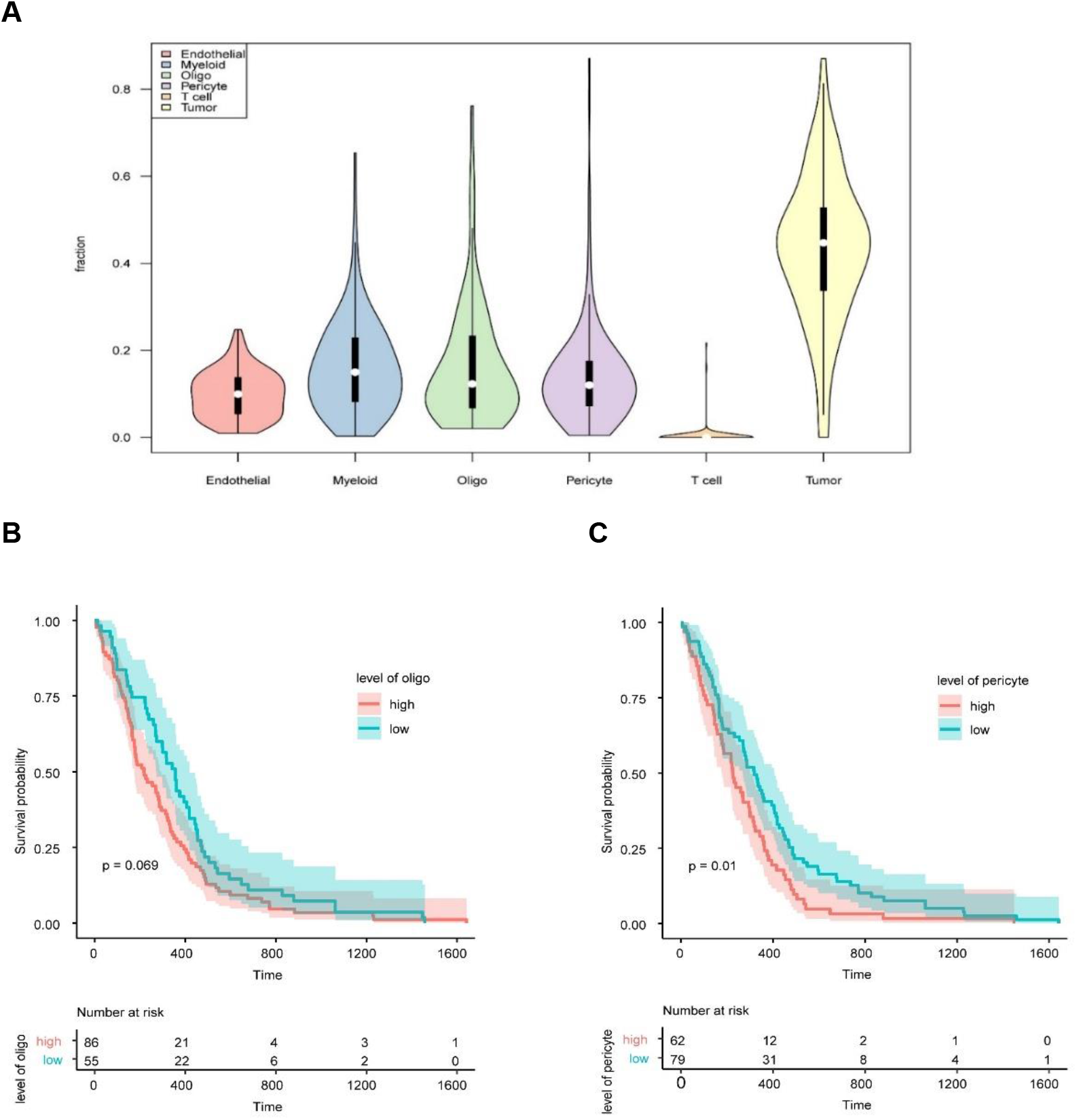
Relationship between cell status and prognosis of non-malignant cells in various tumor types from the TCGA cohort. a, Violin plot visualizing the distribution of cell type fractions in each tumor type. The median is represented by a white dot and the upper and lower quartiles are represented by bars. b,c, The association between oligodendrocyte(b) and pericyte(c) infiltration with survival in GBM using Kaplan-Meier plots.

We also investigated whether there was a correlation between non-malignant cell types and patient survival. Since the TCGA cohort samples varied substantially with respect to treatment, genetic drivers, and other confounding factors, we excluded samples with genetic or clinical covariates, such as IDH-mutant GBMs, which have well-documented and substantial effects on prognosis. To assess the relationship between cell type abundance and patient survival, we utilized two Cox proportional hazards models. To stratify the samples based on cell type abundance, we utilized the cutoff package (Budczies et al. 2012) which allowed us to classify the samples into high and low abundance groups. To assess the relationship between cell type and patient survival, we used cell type abundance as a variable value in our analysis.

Our analysis revealed some important relationships between various immune cell types and clinical outcomes. Our study of GBM revealed that oligodendrocytes(Oligo) and pericyte cells exhibited a stronger correlation with patient survival rate (Fig. 6b,c), which is consistent with previous reports (He et al. 2001; Zhou et al. 2017; Guerra et al. 2018; Hide et al. 2018). The results of our study emphasize the significance of comprehensively understanding the TME of GBM and may have implications for the development of novel therapeutic strategies targeting non-malignant cell types. Overall, our study demonstrates the utility of SCROAM in analyzing bulk RNA-seq to unravel the complex cellular composition of tumors and identify potential prognostic biomarkers.

## Discussion and Conclusion

Accurate identification of cellular composition in complex tissues is crucial for understanding disease pathogenesis, early diagnosis, and prevention. The high cost and technical noise associated with scRNA-seq can limit its widespread use, despite its ability to offer valuable insights into cellular composition. Consequently, cost-effective alternatives to deconvolution of bulk RNA-seq are necessary.

However, most statistical and computational methods have been developed for cell-type deconvolution of bulk RNA-seq, many of these methods have limitations. For example, some methods necessitate prior knowledge of the gene expression profile or cell type composition of purified cell types. There are also methods that rely on a pre-selected list of marker genes. Moreover, fully unsupervised deconvolution methods based on non-negative matrix factorization may suffer from several limitations, such as low precision, discriminability, and multicollinearity issues. Although cell type decomposition on bulk RNA-seq data using scRNA-seq references is an appealing analysis strategy, it is important to note that there may be significant differences in data distribution between scRNA-seq and bulk RNA-seq data, and unreliable feature matrices generated from single-cell references can lead to the identification of incorrect cell types.

To address these challenges and make better use of existing data, we propose SCROAM, a method that utilizes single-cell references and domain-adaptive matching to deconvolute bulk RNA-seq data. By utilizing scRNA-seq references to create reference cell type expression profiles, we eliminate the need for marker gene selection, thereby improving computational efficiency. We calculate the similarity between the true and ideal expression of each gene in the cellular dimension as its specific contribution. Additionally, by reducing the dimensionality of scRNA-seq and bulk RNA-seq to a common feature space, we use domain adaptation matching to learn the difference between the two data in the latent space. After continuous iteration, we learn a weight that eliminates the latent space difference. When significant technical disparities exist between single-cell reference data and observed bulk data, the performance of decomposition can be significantly improved.

To evaluate our method, we benchmarked existing methods using generated pseudo-batch datasets and real experimental datasets with known cell type composition as ground truth. Our results show that SCROAM outperforms existing methods in both settings, providing a more accurate breakdown of cell types. We also applied SCROAM to a mouse spinal cord dataset, where our method successfully identified continuous cell types consistent with biological facts. This analysis further demonstrates the robustness and accuracy of our method in deconvoluting complex tissue samples into their constituent cell types. Finally, we validated our method on liver cancer and tumor datasets and discovered cell types associated with patient prognosis, highlighting the potential of deconvolution to reveal cellular heterogeneity in complex biological systems. Our study demonstrates the versatility of SCROAM and its potential in addressing diverse biological questions of interest. Our approach could provide valuable insights into changes in cell proportions in diseased tissues, informing subsequent therapeutic strategies.

## Materials and Methods

### Data Sources

We evaluated the performance of computational deconvolution using two ways. Firstly, we performed “pseudobulk” experiments, which is a popular approach that involves aggregating scRNA-seq measurements from cells to create gene expression mixtures proportional to the known cell types. By utilizing pseudobulk experiments, we were able to simulate bulk RNA-seq data and evaluate the accuracy of our deconvolution algorithm for diverse cell types. Secondly, we utilized bulk RNA-seq with known cell types proportions to further evaluate the performance of our deconvolution algorithm. This approach allowed us to compare our results with the known cell type proportions, providing an additional measure of the accuracy of our method. We benchmarked our method using these two approaches.

#### Pseudobulk data

In our study, we utilized data from The Tabula Muris Consortium (2020) which includes data on numerous organs/tissues and many cell types through two distinct single-cell experimental protocols: Smart-seq2 (Picelli et al. 2014) and 10x Genomics Chromium. For our in silico experiments, we specifically selected on the following organs/tissues: Kidney, Large_intestine, Liver, Lung, Pancreas, Skin, Thymus, and Trachea (as shown in table 2) for our in silico experiments. We performed two different deconvolutions for each tissue. To perform cross-protocol deconvolution, in one configuration, we used 10x Genomics Chromium as single-cell reference and Smart-seq2 as the pseudobulk, and vice versa for the other configuration.

**Table 2.**
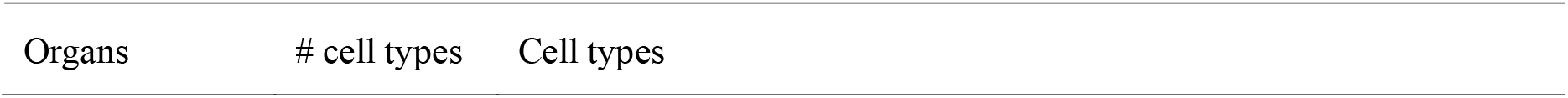
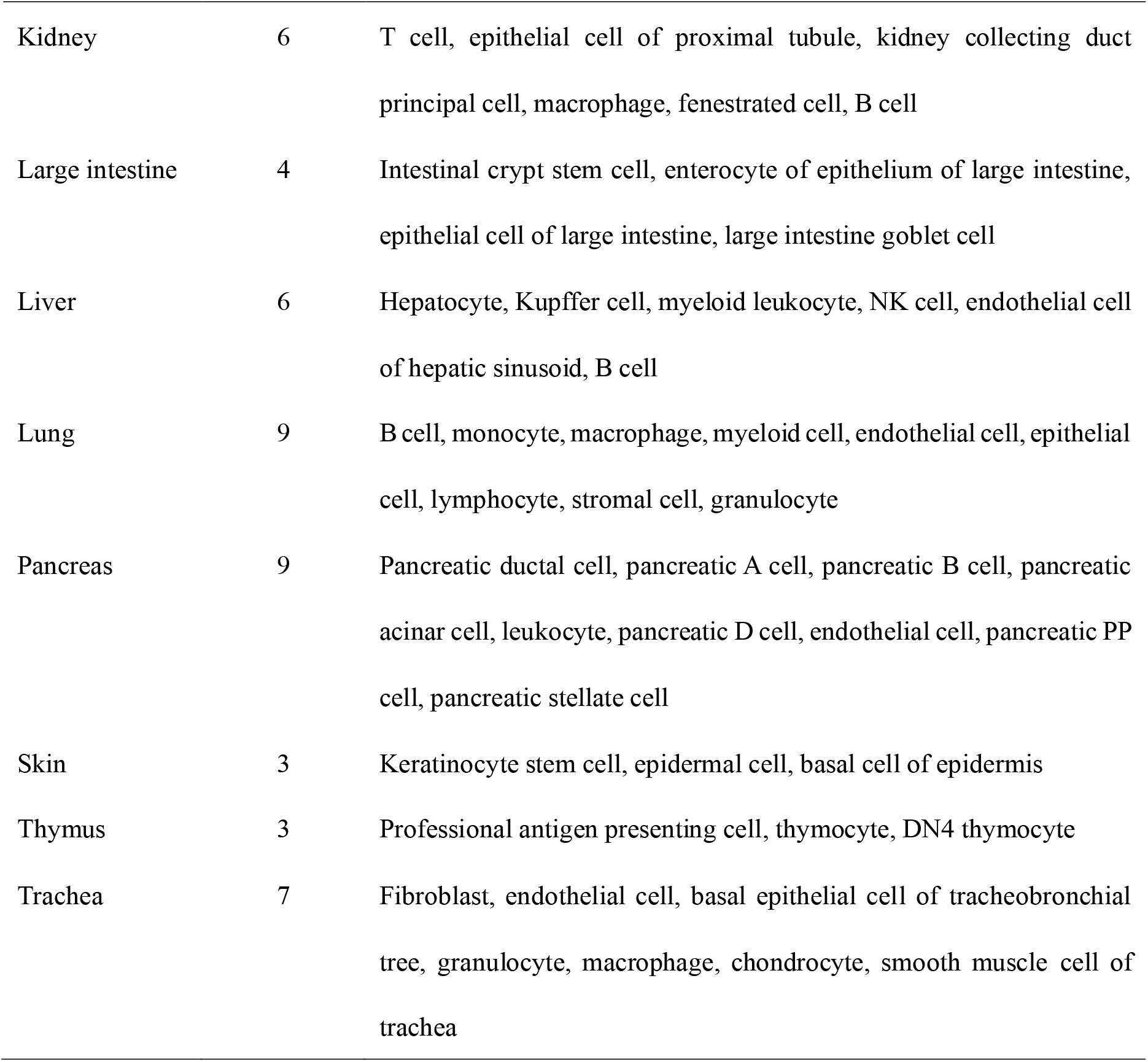
Cell types for every organ in the pseduobulk experiments.

| Organs | # cell types | Cell types |
| --- | --- | --- |
| Kidney | 6 | T cell, epithelial cell of proximal tubule, kidney collecting duct principal cell, macrophage, fenestrated cell, B cell |
| Large intestine | 4 | Intestinal crypt stem cell, enterocyte of epithelium of large intestine, epithelial cell of large intestine, large intestine goblet cell |
| Liver | 6 | Hepatocyte, Kupffer cell, myeloid leukocyte, NK cell, endothelial cell of hepatic sinusoid, B cell |
| Lung | 9 | B cell, monocyte, macrophage, myeloid cell, endothelial cell, epithelial cell, lymphocyte, stromal cell, granulocyte |
| Pancreas | 9 | Pancreatic ductal cell, pancreatic A cell, pancreatic B cell, pancreatic acinar cell, leukocyte, pancreatic D cell, endothelial cell, pancreatic PP cell, pancreatic stellate cell |
| Skin | 3 | Keratinocyte stem cell, epidermal cell, basal cell of epidermis |
| Thymus | 3 | Professional antigen presenting cell, thymocyte, DN4 thymocyte |
| Trachea | 7 | Fibroblast, endothelial cell, basal epithelial cell of tracheobronchial tree, granulocyte, macrophage, chondrocyte, smooth muscle cell of trachea |

The order in which the cell types are listed here corresponds to the order in any figures that were based on our pseudobulk experiments using the Tabula Muris Senis data.

#### Real bulk RNA-seq data

In this section, we offer a detailed description of the real datasets that were used to compare different methods. The first dataset was obtained from Dong(Dong et al. 2021) and consists of human breast cancer and fibroblast cell lines and mixtures. In the second dataset, we obtained bulk whole blood samples from Newman (Newman et al. 2019). However, neutrophils were not present in the single-cell references matched to the bulk RNA-seq, so we used neutrophils from Xie(Xie et al. 2020) instead. The third dataset was obtained from Monaco (Monaco et al. 2019) and consists of bulk whole blood, with single-cell references from 10x Genomics and Xie(Xie et al. 2020) used for comparison. The final dataset used for method comparison was generated from our own biological experiment on mouse spinal cord tissue, with both bulk and single-cell RNA sequencing data obtained. The details of the methods used to generate and analyze the data are provided in the subsequent sections.

##### Spinal Cord-Derived NSCs Culture and Single Cell Solution Preparation

Animal experiments in this study were conducted in strict accordance with the 1996 revised National Institutes of Health Guide for the Care and Use of Laboratory Animals (NIH Publication No. 80-23) and were approved by the Medical Ethics Committee of Xijing Hospital of the Fourth Military Medical University. We made every effort to minimize animal suffering and reduce the number of animals used.

To isolate neural stem cells (NSCs), we followed a previously published method(Zhao et al. 2021) and extracted cells from the spinal cords of 6-week-old adult C57/BL6 mice. In brief, we used three female adult C57/BL6 mice for our experiments. The mice were anesthetized with 5% isoflurane and underwent laminectomy at the T6-12 vertebral level. The T6-T12 spinal cords were isolated from the mice and placed in 10 cm Petri dishes containing cold PBS. The tissue was then dissociated and filtered through 70 µm filters (BD Falcon) to obtain a single-cell suspension. The isolated cells were seeded in NSC growth medium consisting of Dulbecco’s Modified Eagle Medium (DMEM/F12), B-27, and N-2 supplements (Gibco) at a density of 1 × 106 cells per ml.

To prepare single-cell solutions, we subjected the NSCs to enzymatic digestion with Accutase (Gibco) for two passages of 10 minutes at 37°C. The enzymatic digestion was then stopped with NSC growth medium, and the resulting cell suspension was filtered through a 70 μm filter (BD Falcon). The single-cell solutions were stored on ice and loaded into BD Rhapsody cassettes to capture the single-cell transcriptomes.

##### Single cell transcriptome capturation, Library construction and sequencing

In order to preserve cell precision and viability, we used a staining protocol involving two fluorescent materials, Calcein AM and Draq7, and analyzed the cells using a BD Rhapsody™ Scanner (BD Biosciences). The cells were then loaded into a BD Rhapsody microwell cassette according to the protocol developed by Fan (Fan et al. 2015). To ensure that nearly every microwell contained a single cell capture bead, we loaded an excess of beads and washed off any excess beads from the column. Once the cells have been lysed with the lysis buffer, the cell capture beads need to be recovered and washed before reverse transcription can be performed. To generate bead-captured single-cell transcriptomes containing cell label and unique molecular identifier (UMI) information, we used the BD Rhapsody cDNA Kit (BD Biosciences, Cat. No. 633773) and the BD Rhapsody Targeted mRNA and AbSeq Amplification Kit (BD Biosciences, Cat. No. 633801) following the manufacturer’s protocol. All libraries were sequenced in paired-end 150bp mode using the NovaSeq platform (Illumina).

##### Sequencing data processing

We processed the raw sequencing reads from a cDNA library using the BD Rhapsody Whole Transcriptome Assay Analysis Pipeline (v1.8). The pipeline includes several steps, such as quality filtering, annotation, identification of cells, and generation of single-cell expression matrices. The output file contains a matrix of UMI counts per cell and gene, which was used for downstream analysis.

##### Dimensionality reduction, clustering and visualization

First, we used the Seurat R package(Butler et al. 2018; Stuart et al. 2019) to read the original counts matrix of each sample and create a Seurat object. We then filtered out poor quality cells based on the following parameters: percentage of mitochondria below 20, number of UMIs per cell greater than 500, and number of detected genes greater than 200. We performed normalization and scaling, followed by PCA dimensionality reduction based on the top 2000 highly variable genes. Clustering was performed using the Seurat package’s built-in clustering method. To visualize the dimensionality reduction and clustering results, we used t-SNE.

To benchmark SCROAM against other available methods, including Bisque (Jew et al. 2020), MuSiC (Wang et al. 2019), SCDC (Dong et al. 2021) and Non-Negative Least Squares (NNLS) (Mullen 2007), we used all algorithms in each deconvolution with the same data. We ran all algorithms using their respective tutorials with default settings, unless otherwise specified. The NNLS results were obtained from MuSiC’s implementation.

## Methods

### Bulk RNA-seq data transformation

Inspired by domain adaptation learning, deep learning techniques can be utilized to address the problem of sequencing data differences by treating two types of sequencing data as the training set and test set of samples. Our approach to eliminate technical bias between different sequencing data involves the following steps: for a given sequencing data, we obtain its corresponding feature space and calculate the difference between the two feature spaces as the weight of the sample. These weights are then used to continuously adjust the classifier to eliminate any differences in data distribution. We adopt the Kernel Mean Matching Algorithm (Gretton et al. 2009) strategy from the work of Gretton, which is robust to noise and outliers and does not make any assumptions about the sample data.

To begin, we first intersect the genes between scRNA-seq and bulk RNA-seq data to obtain the public gene G. The dimensionality of each cell or sample was then defined as the number of common genes, which was denoted as d. In d-dimensional space, we obtain a set 𝑥_𝑏𝑢𝑙𝑘_ = {𝑥_𝑖_, 𝑖 = 1, …, 𝑛_𝑏𝑢𝑙𝑘_} consisting of 𝑛_𝑏𝑢𝑙𝑘_ independent and identically distributed (i.i.d) samples expressions with a probability density function 𝑝_𝑏𝑢𝑙𝑘_(𝑥), as well as another set 𝑥_𝑠𝑐_ = {𝑥_𝑗_, 𝑗 = 1, …, 𝑛_𝑠𝑐_} consisting of 𝑛_𝑠𝑐_ independent and identically distributed (i.i.d) cells from a different distribution with a probability density function 𝑝_𝑠𝑐_(𝑥). To make the different sample distributions consistent, we estimate the density ratio 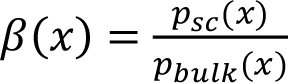 from the given finite samples 𝑥_𝑏𝑢𝑙𝑘_ and 𝑥_𝑠𝑐_.

Next, we transform them from its original high-dimensional space to a Reproducing Kernel Hilbert Space (RKHS) by mapping 𝛷(𝑥): 𝑥 → ℱ. We aim to minimize the weighted distribution 𝛽(𝑥)𝑝_𝑏𝑢𝑙𝑘_(𝑥) and the distribution 𝑝_𝑠𝑐_(𝑥) in Maximum Mean Discrepancy (MMD) (Gretton et al. 2006) between the RKHS to estimate sample weights

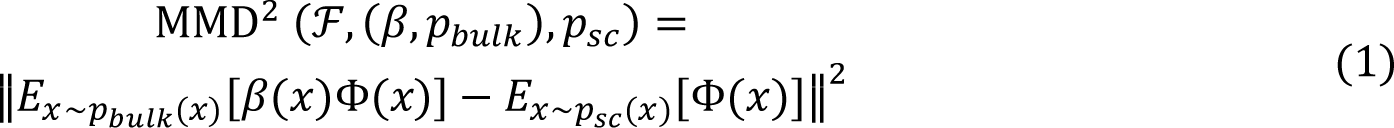

Using the average value of 𝑥_𝑏𝑢𝑙𝑘_ and 𝑥_𝑠𝑐_ instead of the expected value, the new equation is:

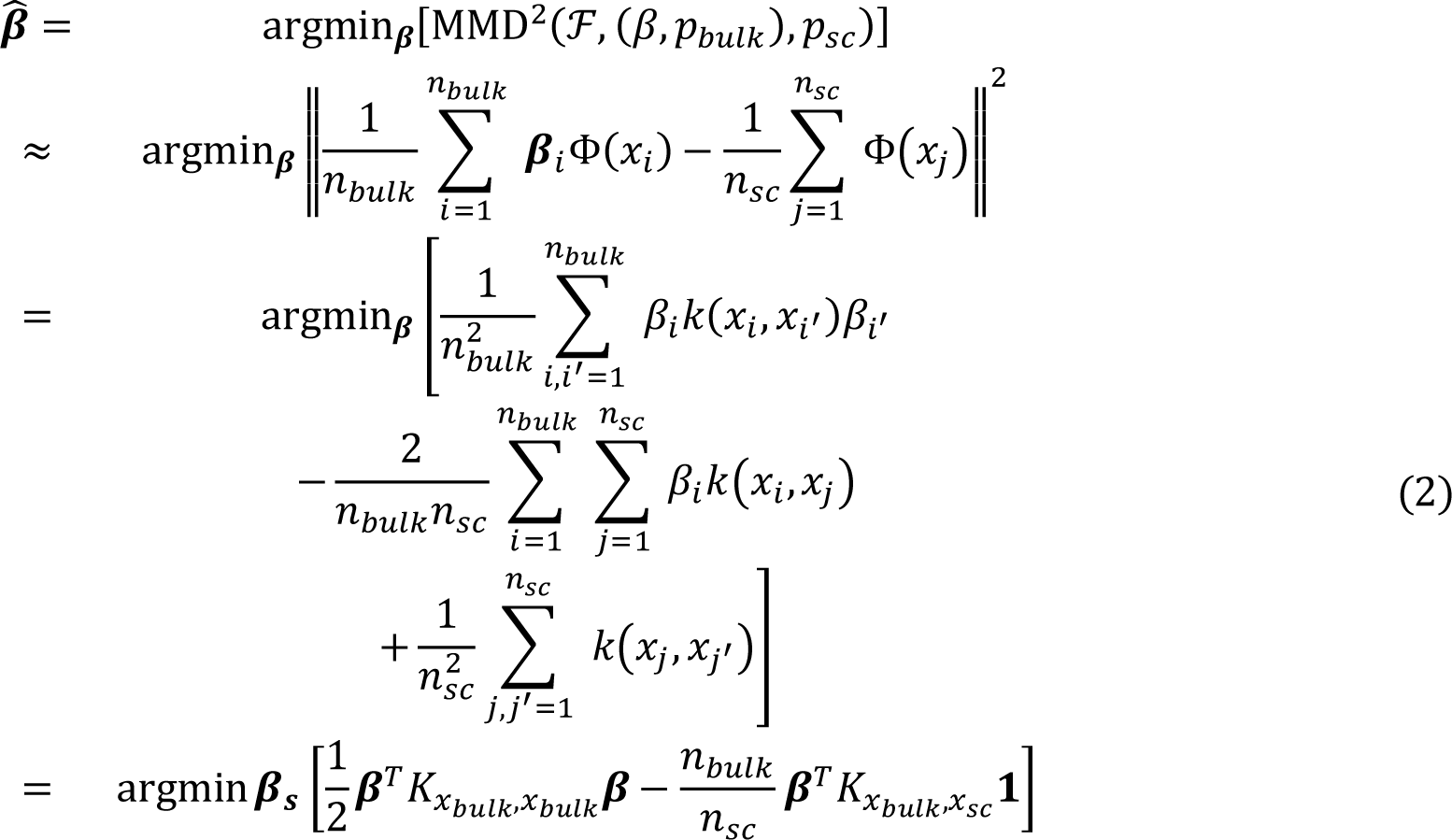

Among them, 𝐾_𝑥𝑏𝑢𝑙𝑘,𝑥𝑏𝑢𝑙𝑘_ and 𝐾_𝑥𝑏𝑢𝑙𝑘,𝑥𝑠𝑐_ are two Gram kernel matrices, and 𝟏_𝑛𝑠𝑐,1_ is a vector of size 𝑛_𝑠𝑐_ × 1.

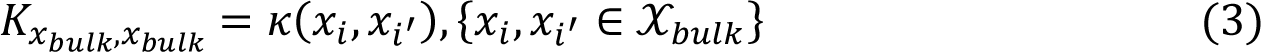

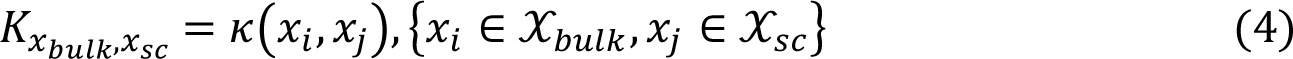

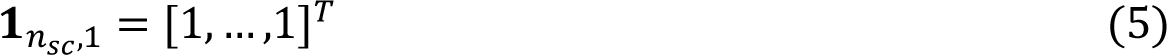

This new equation is subject to two constraints. The first constraint bounds 𝛽_𝑖_ between 0 and B, reflecting the range of differences between 𝑝_𝑏𝑢𝑙𝑘_(𝑥) and 𝑝_𝑠𝑐_(𝑥). The second constraint 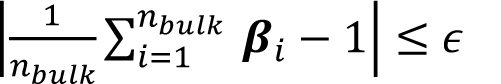 is the normalization of 𝛽(𝑥), because 𝑝_𝑠𝑐_ (𝑥) = 𝛽(𝑥)𝑝_𝑏𝑢𝑙𝑘_ (𝑥) should approximate the probability density function. Small values of ε represent normalized precision. We set the parameter B to the default value of 1000, 𝜖 to 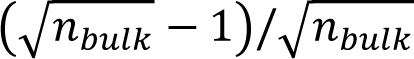 and the kernel bandwidth σ to the median of pairwise sample distances(Gretton et al. 2006), which more detailed information on parameter selection can be found in (Miao et al. 2013). Solving Equation 2 requires solving a convex quadratic programming (QP) problem with linear constraints, which can be accomplished using any existing QP solver (Boyd and Vandenberghe 2004).

#### Construct cell type signature matrix

Single-cell transcriptome data were used to construct feature matrices for cell type deconvolution. Regression-based deconvolution methods typically formulate the deconvolution problem as a system of linear equations, 𝑆𝑥 = 𝑡, where S is a 𝑛 × 𝑘 gene feature matrix (n is the number of genes, k is the number of cell types), and t is a vector of 𝑛 × 1, representing the bulk RNA-seq data. The vector x is a 𝑘 × 1 vector that represents the proportion of cell types, where 𝑛 ≫ 𝑘, resulting in an overdetermined equation and has no exact solution. Most existing methods for solving the overdetermined equation 𝑆𝑥 = 𝑡 are based on the ordinary least squares (OLS) method, which involves minimizing the sum of squares of the absolute error(Abbas et al. 2009; Gong et al. 2011; Gong and Szustakowski 2013; Zhong et al. 2013; Li et al. 2016). However, this approach can result in undesirable consequences, as it may not effectively take into account all informative genes. For instance, if the gene has a low average expression level, its contribution may be small even if its expression varies widely between various cell types. Therefore, we propose the new approach for constructing a feature matrix, inspired by the single-cell differential marker gene approach (Dai et al. 2022). In this strategy, cell types are identified using cluster analysis, and the specificity contribution of each gene is calculated by the cosine similarity between genes for each cell type, and then averaging the gene expression levels for each gene expressed in all cells associated with the cell type are then averaged and multiplied with the contribution matrix, resulting in the cell type signature matrix S. The details are as follows.

To compare the expression patterns of two genes in a given cell cluster, we can represent the expression of each gene using vectors in an n-dimensional cell space. In this space, we create a vector for each gene, where each vector consists of n bases. Here, n represents the total number of detected cells in the population. The value of the vector is the expression level of a gene in all cells. After normalization, we obtain the gene expression matrix 𝑋 ∈ 𝑅^𝑁×𝑀^, where N is the number of cells and M is the number of genes. For the 𝑖_𝑡ℎ_ gene, 𝑔_𝑖_ expressed in all cells is represented by the 𝑖_𝑡ℎ_ column of 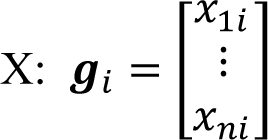, where 𝑋_𝑗𝑖_ is the expression of 𝑔_𝑖_ of the 𝑗_𝑡ℎ_ cell 𝑐_𝑗_, 𝑗 ∈ {1, …, 𝑁}. The variable K is ^𝑥^𝑛𝑖 typically used to denote the number of cell classes, which are either predefined through manual annotation or identified through unsupervised cell clustering. To create an ideal gene 𝑣_𝑘_ that is expressed only in a given cell type 𝐶_𝑘_, 𝑘 ∈ {1, …, 𝐾} and not in any other cell population, we set the ^𝑣^1𝑘 ideal marker gene 𝑣_𝑘_ in 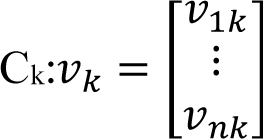. Here, 𝑣_𝑗𝑘_ = 1 if 𝑐_𝑗_ ∈ 𝐶_𝑘_ and 𝑣_𝑗𝑘_ = 0 if 𝑐_𝑗_ ∉ 𝐶_𝑘_ ^𝑣^𝑛𝑘

The cosine similarity between two genes in a scRNA-seq dataset is calculated as the cosine of the angle between their representation vectors in the cell space. A smaller angle indicates that the expression patterns of the two genes are more similar. If the expression patterns of the two genes are identical, the angle between their representative vectors will be zero, independent of their differences in abundance. To compare the representation vector (𝑔_𝑖_, 𝑖 ∈ {1, …, 𝐺}) of each gene with the ideal representation vector 𝑣_𝑘_, we calculate the cosine similarity between 𝑔_𝑖_ and 𝑣_𝑘_, using the formula 𝑐𝑜𝑠(𝑔_𝑖_, 𝑣_𝑘_)

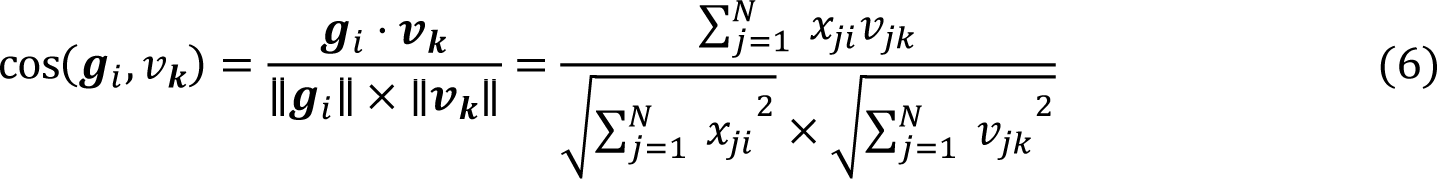

To determine the gene-specific score of each gene in each cell type, we aim to maximize the cosine similarity between the gene’s representation vector and the ideal vector 𝑣_𝑘_ of its corresponding cell type while minimizing the cosine similarity between the gene’s representation vector and the ideal vector of other cell types. We calculate the 𝑐𝑜𝑛𝑡𝑟𝑖𝑏𝑢𝑡𝑒_𝑠𝑐𝑜𝑟𝑒(𝑔𝑖,𝐶𝑘)_ for each gene as follows:

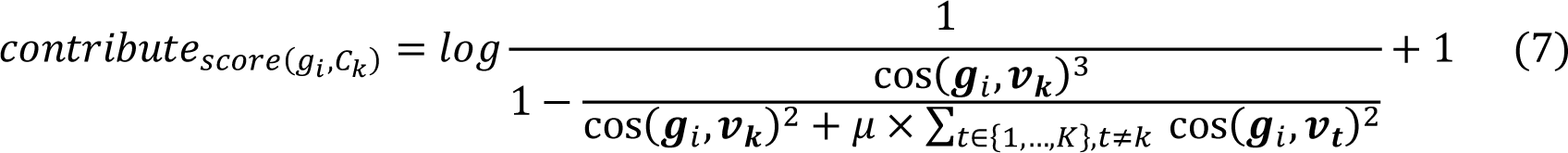

Here, 𝑢 ≥ 0 represents the degree of penalty imposed on a gene’s representation vector when it matches the ideal vector of a non-target cell type (𝐶_𝑡_, 𝑡 ∈ {1, …, 𝐾} and 𝑡 ≠ 𝑘). The value of u can be adjusted by the user, with a higher value of u indicating a greater punishment for genes that are expressed in non-target cell types. By default, we set u=1.

#### Cell type proportion estimation

This section describe the connection between bulk and scRNA-seq gene expression. This relationship is essential to our deconvolution process, which we estimate cell type proportions using the W-NNLS framework proposed by MuSiC (Wang et al. 2019). After normalization, 𝑋_𝑔𝑐_ represents all the expression levels of gene g in cell c, and the total number of cells in cell type k is denoted as 𝑚^𝑘^ = |𝐶^𝑘^|. Thus, the total abundance of gene g is given by 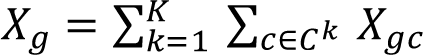 and the total number of cells is denoted as 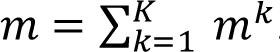. The fraction of cells from cell type k is given by 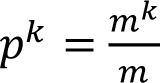. We can 𝑚 calculate the average abundance of gene g in cell type k using the following formula:

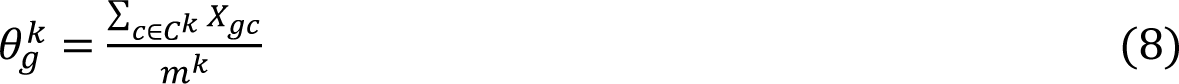

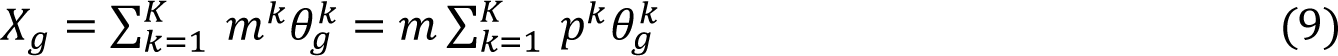

The relative abundance of gene g in bulk tissue can be expressed as 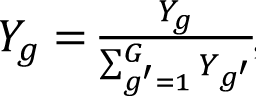, where Y = 𝛽𝑥_bulk_ is the transformed gene expression in bulk tissues, and G is the total number of genes. Therefore,

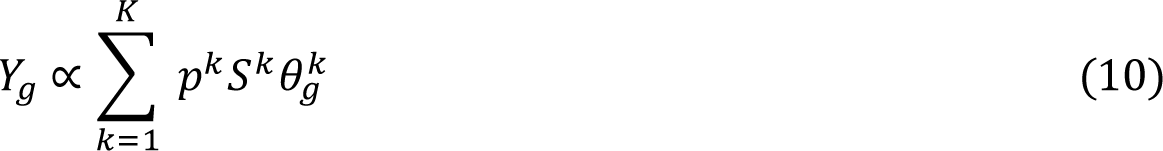

S is the feature matrix we calculated and is a constant.

To estimate 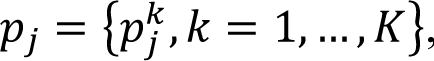, We need to satisfy two constraints. The first constraint is non-negativity, which requires that 𝑝^𝑘^_*j*_ ≥ 0,for all j and k. The second constraint is 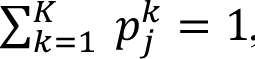, for all j. Since the relationship between bulk and scRNA-seq expression profiles, as deduced in formula (10), is a “proportional” relationship, we need a normalization constant to satisfy the second constraint.

## Data sets

All the data used in this study are publicly available and can be accessed through the accession numbers provided in Supplemental Table S3. The self-sequenced data has been uploaded to GitHub and can be accessed through the Genome Research website.

## Acknowledgments

This work was supported by the National Natural Science Foundation of China (Grants #62072353 and #62272065 and #62271049).

## Author Contributions

L.Y. and C.G.Z. initiated and envisioned the study. X.Y.G, Z.Y.H. and L.Y. formulated the model. X.Y.G. was responsible for implementing the algorithm and performing simulation studies in this research. Z.Y.H and F.J. performed a baseline comparison of human breast cancer and peripheral blood. C.G.Z. and F.J. conceived and carried out the biological experiments. All authors participated in the real data analysis presented in this paper. X.Y.G. and L.Y. were responsible for writing the manuscript, which was subsequently reviewed, edited, and approved by all authors.

## References

1. Abbas AR, Wolslegel K, Seshasayee D, Modrusan Z, Clark HF. 2009. Deconvolution of blood microarray data identifies cellular activation patterns in systemic lupus erythematosus. PloS one 4: e6098. doi: 10.1371/journal.pone.0006098

2. Ally A, Balasundaram M, Carlsen R, Chuah E, Clarke A, Dhalla N, Holt RA, Jones SJ, Lee D, Ma Y. 2017. Comprehensive and integrative genomic characterization of hepatocellular carcinoma. Cell 169: 1327–1341. e1323. doi: 10.1016/j.cell.2017.05.046

3. Aran D, Hu Z, Butte AJ. 2017. xCell: digitally portraying the tissue cellular heterogeneity landscape. Genome Biol 18: 1–14. doi: 10.1186/s13059-017-1349-1

4. Avila Cobos F, Vandesompele J, Mestdagh P, De Preter K. 2018. Computational deconvolution of transcriptomics data from mixed cell populations. Bioinformatics 34: 1969–1979. doi: 10.1093/bioinformatics/bty019

5. Becht E, Giraldo NA, Lacroix L, Buttard B, Elarouci N, Petitprez F, Selves J, Laurent-Puig P, Sautès-Fridman C, Fridman WH. 2016. Estimating the population abundance of tissue-infiltrating immune and stromal cell populations using gene expression. Genome Biol 17: 1–20. doi: 10.1186/s13059-016-1070-5.

6. Boyd SP, Vandenberghe L. 2004. Convex optimization. Cambridge university press.

7. Brennan CW, Verhaak RG, McKenna A, Campos B, Noushmehr H, Salama SR, Zheng S, Chakravarty D, Sanborn JZ, Berman SH, et al. 2013. The somatic genomic landscape of glioblastoma. Cell 155: 462–477. doi: 10.1016/j.cell.2013.09.034.

8. Budczies J, Klauschen F, Sinn BV, Győrffy B, Schmitt WD, Darb-Esfahani S, Denkert C. 2012. Cutoff Finder: a comprehensive and straightforward Web application enabling rapid biomarker cutoff optimization. PloS one 7: e51862. doi: 10.1371/journal.pone.0051862

9. Butler A, Hoffman P, Smibert P, Papalexi E, Satija R. 2018. Integrating single-cell transcriptomic data across different conditions, technologies, and species. Nat Biotechnol 36: 411–420. doi: 10.1038/nbt.4096

10. Carithers LJ, Moore HM. 2015. The genotype-tissue expression (GTEx) project. Biopreserv Biobank 13: 307–308. doi: 10.1089/bio.2015.29031

11. Chen L. 2019. CAMTHC: convex analysis of mixtures for tissue heterogeneity characterization.

12. Chung BK, Karlsen TH, Folseraas T. 2018. Cholangiocytes in the pathogenesis of primary sclerosing cholangitis and development of cholangiocarcinoma. Biochim Biophys Acta Mol Basis Dis 1864: 1390–1400. doi: 10.1016/j.bbadis.2017.08.020

13. Cohen I, Huang Y, Chen J, Benesty J, Benesty J, Chen J, Huang Y, Cohen I. 2009. Pearson correlation coefficient. Noise reduction in speech processing: 1–4. doi: 10.1007/978-3-642-00296-0_5

14. Dai M, Pei X, Wang X-J. 2022. Accurate and fast cell marker gene identification with COSG. Briefings in Bioinform 23: bbab579. doi: 10.1093/bib/bbab579

15. Denisenko E, Guo BB, Jones M, Hou R, De Kock L, Lassmann T, Poppe D, Clément O, Simmons RK, Lister R. 2020. Systematic assessment of tissue dissociation and storage biases in single-cell and single-nucleus RNA-seq workflows. Genome Biol 21: 1–25. doi: 10.1186/s13059-020-02048-6

16. Dong M, Thennavan A, Urrutia E, Li Y, Perou CM, Zou F, Jiang Y. 2021. SCDC: bulk gene expression deconvolution by multiple single-cell RNA sequencing references. Brief Bioinform 22: 416–427. doi: 10.1093/bib/bbz166

17. Erdmann-Pham DD, Fischer J, Hong J, Song YS. 2021. Likelihood-based deconvolution of bulk gene expression data using single-cell references. Genome Res 31: 1794–1806. doi: 10.1101/gr.272344.120

18. Fan HC, Fu GK, Fodor SP. 2015. Expression profiling. Combinatorial labeling of single cells for gene expression cytometry. Science 347: 1258367. doi: 10.1126/science.1258367

19. Finotello F, Trajanoski Z. 2018. Quantifying tumor-infiltrating immune cells from transcriptomics data. Cancer Immunol Immunother 67: 1031–1040. doi: 10.1007/s00262-018-2150-z

20. Gong T, Hartmann N, Kohane IS, Brinkmann V, Staedtler F, Letzkus M, Bongiovanni S, Szustakowski JD. 2011. Optimal deconvolution of transcriptional profiling data using quadratic programming with application to complex clinical blood samples. PloS one 6: e27156. doi: 10.1371/journal.pone.0027156

21. Gong T, Szustakowski JD. 2013. DeconRNASeq: a statistical framework for deconvolution of heterogeneous tissue samples based on mRNA-Seq data. Bioinformatics 29: 1083–1085. doi: 10.1093/bioinformatics/btt090

22. Gracia-Sancho J, Caparrós E, Fernández-Iglesias A, Francés R. 2021. Role of liver sinusoidal endothelial cells in liver diseases. Nat Rev Gastroenterol Hepatol 18: 411–431. doi: 10.1038/s41575-020-00411-3.

23. Gretton A, Borgwardt K, Rasch M, Schölkopf B, Smola A. 2006. A kernel method for the two-sample-problem. Advances in neural information processing systems 19. doi: 10.48550/arXiv.0805.2368

24. Gretton A, Smola A, Huang J, Schmittfull M, Borgwardt K, Schölkopf B. 2009. Covariate shift by kernel mean matching. Dataset shift in machine learning 3: 131–160. doi: 10.7551/mitpress/9780262170055.003.0008

25. Guerra DA, Paiva AE, Sena IF, Azevedo PO, Silva WN, Mintz A, Birbrair A. 2018. Targeting glioblastoma-derived pericytes improves chemotherapeutic outcome. Angiogenesis 21: 667–675. doi: 10.1007/s10456-018-9621-x

26. He J, Mokhtari K, Sanson M, Marie Y, Kujas M, Huguet S, Leuraud P, Capelle L, Delattre J, Poirier J. 2001. Glioblastomas with an oligodendroglial component: a pathological and molecular study. J Neuropathol Exp Neurol 60: 863–871. doi: 10.1093/jnen/60.9.863

27. Hide T, Komohara Y, Miyasato Y, Nakamura H, Makino K, Takeya M, Kuratsu J-i, Mukasa A, Yano S. 2018. Oligodendrocyte progenitor cells and macrophages/microglia produce glioma stem cell niches at the tumor border. EBioMedicine 30: 94–104. doi: 10.1016/j.ebiom.2018.02.024

28. Jew B, Alvarez M, Rahmani E, Miao Z, Ko A, Garske KM, Sul JH, Pietiläinen KH, Pajukanta P, Halperin E. 2020. Accurate estimation of cell composition in bulk expression through robust integration of single-cell information. Nature Commun 11: 1971. doi: 10.1038/s41467-020-15816-6

29. Jin H, Liu Z. 2021. A benchmark for RNA-seq deconvolution analysis under dynamic testing environments. Genome Biol 22: 1–23. doi: 10.1186/s13059-021-02290-6

30. Kuksin M, Morel D, Aglave M, Danlos F-X, Marabelle A, Zinovyev A, Gautheret D, Verlingue L. 2021. Applications of single-cell and bulk RNA sequencing in onco-immunology. Eur J Cancer 149: 193–210. doi: 10.1016/j.ejca.2021.03.005

31. Li B, Severson E, Pignon J-C, Zhao H, Li T, Novak J, Jiang P, Shen H, Aster JC, Rodig S. 2016. Comprehensive analyses of tumor immunity: implications for cancer immunotherapy. Genome Biol 17: 1–16. doi: 10.1186/s13059-016-1028-7

32. Liebner DA, Huang K, Parvin JD. 2014. MMAD: microarray microdissection with analysis of differences is a computational tool for deconvoluting cell type-specific contributions from tissue samples. Bioinformatics 30: 682–689. doi: 10.1093/bioinformatics/btt566

33. Macosko EZ, Basu A, Satija R, Nemesh J, Shekhar K, Goldman M, Tirosh I, Bialas AR, Kamitaki N, Martersteck EM. 2015. Highly parallel genome-wide expression profiling of individual cells using nanoliter droplets. Cell 161: 1202–1214. doi: 10.1016/j.cell.2015.05.002

34. MacParland SA, Liu JC, Ma X-Z, Innes BT, Bartczak AM, Gage BK, Manuel J, Khuu N, Echeverri J, Linares I. 2018. Single cell RNA sequencing of human liver reveals distinct intrahepatic macrophage populations. Nature Commun 9: 4383. doi: 10.1038/s41467-018-06318-7

35. Massalha H, Bahar Halpern K, Abu-Gazala S, Jana T, Massasa EE, Moor AE, Buchauer L, Rozenberg M, Pikarsky E, Amit I. 2020. A single cell atlas of the human liver tumor microenvironment. Mol Syst Biol 16: e9682. doi: 10.15252/msb.20209682

36. Menden K, Marouf M, Oller S, Dalmia A, Magruder DS, Kloiber K, Heutink P, Bonn S. 2020. Deep learning–based cell composition analysis from tissue expression profiles. Science Adv 6: eaba2619. doi: 10.1126/sciadv.aba2619

37. Menéndez M, Pardo J, Pardo L, Pardo M. 1997. The jensen-shannon divergence. Journal of the Franklin Institute 334: 307–318

38. Miao Y-Q, Farahat AK, Kamel MS. 2013. Auto-tuning kernel mean matching. In 2013 IEEE 13th International Conference on Data Mining Workshops, pp. 560–567. IEEE. doi: 10.1145/2740908.2742476

39. Monaco G, Lee B, Xu W, Mustafah S, Hwang YY, Carré C, Burdin N, Visan L, Ceccarelli M, Poidinger M. 2019. RNA-Seq signatures normalized by mRNA abundance allow absolute deconvolution of human immune cell types. Cell Rep 26: 1627–1640. e1627. doi: 10.1016/j.celrep.2019.01.041

40. Mullen KM. 2007. nnls: The Lawson-Hanson NNLS algorithm for non-negative least squares.

41. Newman AM, Steen CB, Liu CL, Gentles AJ, Chaudhuri AA, Scherer F, Khodadoust MS, Esfahani MS, Luca BA, Steiner D. 2019. Determining cell type abundance and expression from bulk tissues with digital cytometry. Nat Biotechnol 37: 773–782. doi: 10.1038/s41587-019-0114-2

42. Picelli S, Faridani OR, Björklund ÅK, Winberg G, Sagasser S, Sandberg R. 2014. Full-length RNA-seq from single cells using Smart-seq2. Nat Protoc 9: 171–181. doi: 10.1038/nprot.2014.006

43. Pinto C, Giordano DM, Maroni L, Marzioni M. 2018. Role of inflammation and proinflammatory cytokines in cholangiocyte pathophysiology. Biochim Biophys Acta Mol Basis Dis 1864: 1270–1278. doi: 10.1016/j.bbadis.2017.07.024

44. Poisson J, Lemoinne S, Boulanger C, Durand F, Moreau R, Valla D, Rautou P-E. 2017. Liver sinusoidal endothelial cells: Physiology and role in liver diseases. J Hepatol 66: 212–227. doi: 10.1016/j.jhep.2016.07.009

45. Saliba A-E, Westermann AJ, Gorski SA, Vogel J. 2014. Single-cell RNA-seq: advances and future challenges. Nucleic Acids Res 42: 8845–8860. doi: 10.1093/nar/gku555

46. Stuart T, Butler A, Hoffman P, Hafemeister C, Papalexi E, Mauck WM, Hao Y, Stoeckius M, Smibert P, Satija R. 2019. Comprehensive integration of single-cell data. Cell 177: 1888–1902. e1821. doi: 10.1016/j.cell.2019.05.031

47. Sturm G, Finotello F, Petitprez F, Zhang JD, Baumbach J, Fridman WH, List M, Aneichyk T. 2019. Comprehensive evaluation of transcriptome-based cell-type quantification methods for immuno-oncology. Bioinformatics 35: i436–i445. doi: 10.1093/bioinformatics/btz363

48. Tabula Muris Consortium. 2020. A single-cell transcriptomic atlas characterizes ageing tissues in the mouse. Nature 583: 590–595. doi: 10.1038/s41586-020-2496-1

49. Tomczak K, Czerwińska P, Wiznerowicz M. 2015. Review The Cancer Genome Atlas (TCGA): an immeasurable source of knowledge. Contemp Oncol (Pozn) 2015: 68–77. doi: 10.5114/wo.2014.47136

50. Vallania F, Tam A, Lofgren S, Schaffert S, Azad TD, Bongen E, Haynes W, Alsup M, Alonso M, Davis M. 2018. Leveraging heterogeneity across multiple datasets increases cell-mixture deconvolution accuracy and reduces biological and technical biases. Nat Commun 9: 4735. doi: 10.1038/s41467-018-07242-6

51. Wang N, Gong T, Clarke R, Chen L, Shih I-M, Zhang Z, Levine DA, Xuan J, Wang Y. 2015. UNDO: a Bioconductor R package for unsupervised deconvolution of mixed gene expressions in tumor samples. Bioinformatics 31: 137–139. doi: 10.1093/bioinformatics/btu607

52. Wang X, Park J, Susztak K, Zhang NR, Li M. 2019. Bulk tissue cell type deconvolution with multi-subject single-cell expression reference. Nat Commun 10: 380. doi: 10.1038/s41467-018-08023-x

53. Xie X, Shi Q, Wu P, Zhang X, Kambara H, Su J, Yu H, Park S-Y, Guo R, Ren Q. 2020. Single-cell transcriptome profiling reveals neutrophil heterogeneity in homeostasis and infection. Nat Immunol 21: 1119–1133. doi: 10.1038/s41590-020-0736-z

54. Yuan J, Levitin HM, Frattini V, Bush EC, Boyett DM, Samanamud J, Ceccarelli M, Dovas A, Zanazzi G, Canoll P. 2018. Single-cell transcriptome analysis of lineage diversity in high-grade glioma. Genome Med 10: 1–15. doi: 10.1186/s13073-018-0567-9

55. Zhao C-G, Qin J, Li J, Jiang S, Ju F, Sun W, Ren Z, Ji Y-Q, Wang R, Sun X-L. 2021. LINGO-1 regulates Wnt5a signaling during neural stem and progenitor cell differentiation by modulating miR-15b-3p levels. Stem Cell Res Ther 12: 372. doi: 10.1186/s13287-021-02452-0

56. Zhong Y, Wan Y-W, Pang K, Chow LM, Liu Z. 2013. Digital sorting of complex tissues for cell type-specific gene expression profiles. BMC bioinformatics 14: 1–10. doi: 10.1186/1471-2105-14-89

57. Zhou W, Chen C, Shi Y, Wu Q, Gimple RC, Fang X, Huang Z, Zhai K, Ke SQ, Ping Y-F. 2017. Targeting glioma stem cell-derived pericytes disrupts the blood-tumor barrier and improves chemotherapeutic efficacy. Cell Stem Cell 21: 591–603. e594. doi: 10.1016/j.stem.2017.10.002

